# From cognitive abstraction to adaptive behavior: neural bases of concept learning in autistic adolescents

**DOI:** 10.64898/2026.01.17.700125

**Authors:** Yijun Chen, Brylee Hawkins, Hannah Puckett, Kat Sharp, Andrea Lopez, Dagmar Zeithamova, Hua Xie, Alyssa Verbalis, Ashley S. Van Meter, William D. Gaillard, Lauren Kenworthy, Chandan J. Vaidya

## Abstract

**BACKGROUND:** Learned knowledge does not consistently generalize to new contexts in autistic individuals, limiting potential for adapting to real-world demands. This challenge is hypothesized to stem from difficulties with forming abstract representations, potentially arising from perceptual processing that favors local details over the gestalt. We tested the prediction that generalization would be primarily based on exemplar-specific representations in autistic youth using computational modelling coupled with neuroimaging.

**METHODS:** Sixty-four autistic adolescents without intellectual disability (69% males; ages 14-18 years) completed a category generalization task during functional magnetic resonance imaging at two time points. Computational models estimated abstract (prototype-based) and specific (exemplar-based) representations and underlying neural correlates. We further examined associations with adaptive functioning and moderation by autistic traits.

**RESULTS:** Contrary to predictions, we observed a consistent prototype-dominant majority, a subgroup who generalized without consistent representational reliance, and a small minority who failed to acquire category structure. Prototypes were represented in bilateral ventromedial prefrontal cortex (VMPFC), inferior parietal lobule (IPL), right frontal pole, and right lateral occipital cortex, while exemplars were represented in bilateral cuneus. Better generalization predicted better real-world adaptive functioning. Moreover, greater prototype-related activation in left IPL predicted better adaptive functioning in participants with higher autistic traits.

**CONCLUSIONS:** These findings challenge the prevailing view that concept learning in autism relies primarily on hyper-specific perceptual processing, identify meaningful variability in representational strategies, and reveal neural pathways through which abstract representation may support real-world adaptive behavior.

## Introduction

Autistic individuals do not consistently generalize learned knowledge and skills to novel settings, posing challenges for addressing social, communication, and behavioral symptoms that may limit their functional independence. Weak generalization in autism is posited to reflect difficulties in forming abstract memory representations (De Marchena, Eigsti, & Yerys, 2015; Klinger & Dawson, 2001; Minshew, Meyer, & Goldstein, 2002). Theoretical and computational approaches suggest that abstraction from past experiences promotes transfer to novel contexts, thereby facilitating adaptation to the environment (Wu, Meder, & Schulz, 2025). One form of abstraction involves attending to similarities across experiences and storing a summary representation of a concept, termed *prototype*. Compared with storing many individual experiences, relying on a prototype reduces memory load and provides an efficient way to respond to novel events. It also provides a shorthand for identifying familiar aspects of novel events thereby reducing uncertainty. Such integrative processing may not be automatic for autistic individuals whose perception is reported to favor local details over the gestalt (Happé & Frith, 2006). This perceptual bias is thought to lead to hyper-specific representations (Harris et al., 2015), limiting the potential for generalization. For example, autistic individuals may learn an object category by remembering multiple objects encountered in the past, which may not necessarily help recognize a new object. This view predicts that differences in concept representation underpin generalization difficulty in autism, which in turn, impacts real-world functioning. No study with autistic individuals has examined this prediction comprehensively.

Foundational research on concept learning, typically using category generalization paradigms, has revealed mixed findings about autistic individuals’ potential for abstract processing. Early findings established that autistic children demonstrated learning of real-world object categories, despite intellectual functioning challenges (Tager-Flusberg, 1985a, 1985b; Ungerer & Sigman, 1987) but deficits emerged when categorization required integration across multiple object dimensions (Shulman, Yirmiya, & Greenbaum, 1995). Later studies used artificial stimuli to test whether categorization was based on prototypes. Prototype-based learning is reflected in a “typicality gradient”, with categorization accuracy highest for the prototype and decreasing as stimuli become less similar. Some studies found that overall generalization accuracy and/or typicality effects, were lower in autistic relative to typically developing (TD) individuals (Klinger & Dawson, 2001; Church et al., 2010; Vladusich, Olu-Lafe, Kim, Tager-Flusberg, & Grossberg, 2010; Froehlich et al., 2012; Gastgeb, Dundas, Minshew, & Strauss, 2012; Mercado et al., 2015), while others found no group differences (Molesworth, Bowler, & Hampton, 2005, 2008; Schipul & Just, 2016; Tager-Flusberg, 1985a; Vladusich et al., 2010). Among autistic individuals, two subgroups were noted, one showing typicality deficits and another comparable to TD peers (Church et al., 2010; Molesworth et al., 2008). Age, IQ, or number of training stimuli or categories did not account for the variable results (Vanpaemel & Bayer, 2021), leaving open the question of the extent to which autistic individuals use abstract representation to guide generalization.

Application of computational modelling to estimate representations underlying generalization holds promise for elucidating the observed heterogeneity. Prototype models assume that new stimuli are categorized by comparing similarity to a summary representation of training stimuli (Posner & Keele, 1968), whereas exemplar models assume comparison to individual training stimuli (Nosofsky, 1986). Available evidence comes from only two autistic samples: prototype modelling in 20 children showed that 12 were sensitive to prototype and 8 were not (Church et al., 2010; Voorspoels, Rutten, Bartlema, Tuerlinckx, & Vanpaemel, 2018) and exemplar modelling in 16 adults did not show group differences (Bott, Brock, Brockdorff, Boucher, & Lamberts, 2006). No firm conclusions about representational heterogeneity can be drawn due to different sample ages and their small sizes in these studies. Moreover, no study applied both models in the same autistic individuals to determine which representation best describes generalization (Bowman, Iwashita, & Zeithamova, 2020; Bowman & Zeithamova, 2018). The stability of representational strategies over time should also be considered because hyper-specific perception is posited to be a phenotypic trait rather than state-dependent (Happé & Frith, 2006). Finally, whether generalization and underlying representations predict real-world functioning as theorized (Wu et al., 2025) also remains an open question.

Here we used computational modelling coupled with functional magnetic resonance imaging (fMRI) at two timepoints, nine months apart, to resolve open questions about generalization, abstract representations, and adaptive functioning in autism. Participants were 14–18-year-old autistic youth without intellectual disability. The late adolescent age range is of particular concern in autism because of impending demands to transition to adult independent functioning (White et al., 2021). We tested the following predictions: First, in light of the hypothesized exemplar-specific perceptual bias in autism, we predicted that autistic youth would rely on exemplars over prototypes, consistently over time. Second, based on past model-based fMRI work, we expected prototype representation in ventromedial prefrontal cortex (VMPFC) and anterior hippocampus and exemplar representation in posterior hippocampus (Bowman & Zeithamova, 2018). Finally, considering the importance of generalization and abstract processing for real-world behavior, we expected generalization accuracy and prototype neural correlates to predict parent-reported adaptive functioning.

## Methods and Materials

### Participants

Sixty-four adolescents with diagnosis of autism aged 14–18 years were enrolled for a study with two timepoints of assessments approximately nine months apart (see demographics in Table 1). Autism diagnosis was confirmed based on DSM-5 criteria by an expert clinician, supported by a community-based diagnosis and a Social Communication Questionnaire (SCQ) score ≥7. If diagnosis was not confirmed by the SCQ score, an ADOS was administered or reviewed by the expert clinician. Medication and co-occurring conditions are reported in Supplementary Materials (SM). Exclusion criteria included Wechsler Full Scale IQ score below 80, other neurological diagnosis (e.g., epilepsy) based on parent report, contraindication for MRI, and failing neuroimaging and task performance data quality criteria (described in SM). After data quality controls, the retained sample was N=50 at Timepoint 1 (T1), N=42 at Timepoint 2 (T2) and N=39 at both timepoints for task performance analysis, and N=39 at T1, N=31 at T2; N=27 at both timepoints for neuroimaging analysis. All participants complied with consenting procedures approved by the Institutional Review Boards at Children’s National Hospital and Georgetown University.

**Table 1.**
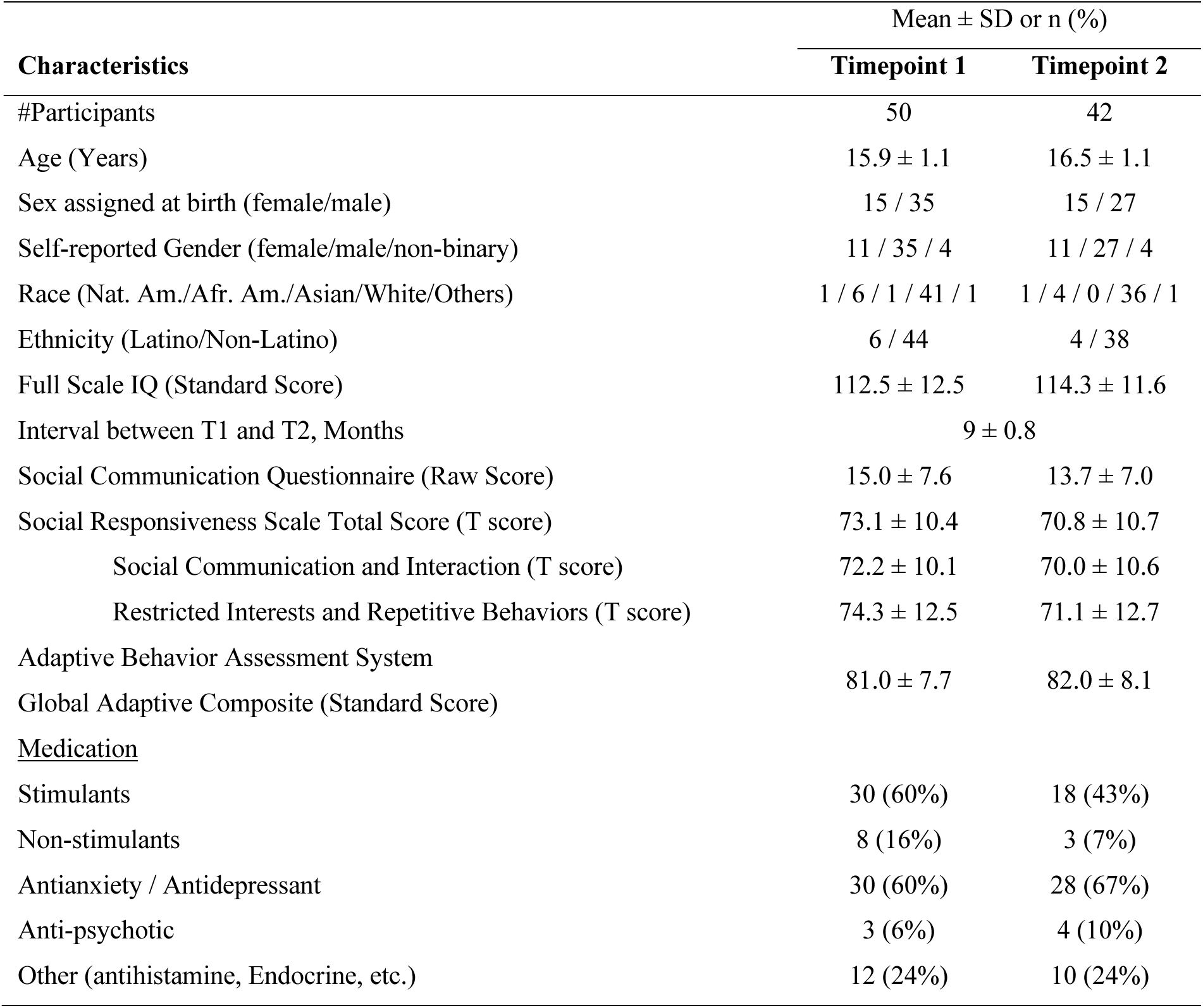
Demographic Characteristics

A parent completed the Adaptive Behavior Assessment System–II Global Adaptive Composite (ABAS-GAC) score for assessing real-world behavior and the Social Responsiveness Scale–II (SRS; see SM for details) for assessing autism traits.

### Category learning task

#### Materials

The category learning task was modelled after Bowman and Zeithamova (Bowman & Zeithamova, 2018) and comprised cartoon animal stimuli sets varying along eight binary features (Figures 1A). For each stimulus set, one exemplar was randomly selected as prototype for Category A and the exemplar with all eight opposing features was the Category B prototype. Thus, the two prototypes were eight features apart. Each category included exemplars that were 1-away, 2-away, and 3-away from their respective prototypes. Exemplars equidistant from the prototypes (i.e., 4-away) were excluded. For each stimulus set, eight 2-away exemplars (four per category) served as training stimuli; the two prototypes and 24 new exemplars per category (eight 1-away, eight 2-away and eight 3-away from their respective prototypes) were used as generalization stimuli, for a total of 58 unique exemplars per set. We designed three stimulus sets (Fish, Bugs, Butterflies; Figure S1) to ensure task novelty across timepoints. Stimulus sets were randomly assigned such that the three sets were represented at each timepoint across participants and each participant received different stimuli at the two timepoints. Each participant was assigned one of four possible prototypes within a set, and left/right category label placement was counterbalanced across participants.

**Figure 1.**
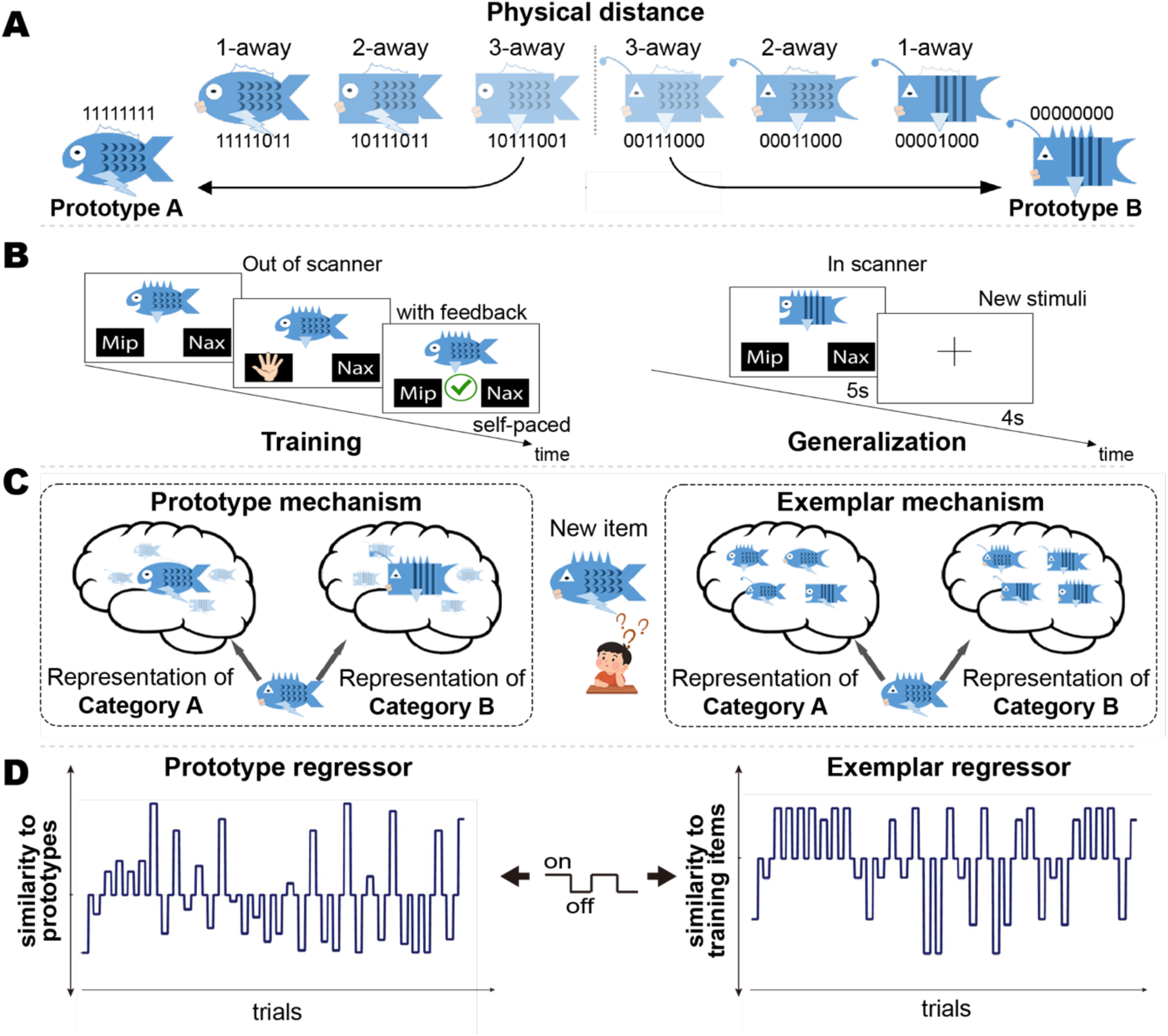
Schematic of experimental design and model-based neural model fitting. (A) Example stimuli spanning *n* feature-distance away (1, 2, and 3) from prototype A and B (0); The two prototypes are 8 features away from each other; thus, a stimulus that is 3-away from one prototype, is 5 features away from the other prototype. (B) Sequence of events in a trial during the training (left) and generalization (right) phases. Training stimuli were 2-feature away and generalization stimuli were 0,1, 2, and 3 features away. (C) Illustrative depictions of category representations as assumed by the prototype and exemplar models. The prototype model assumes that categories are represented by abstract category information combining all typical category features into an idealized prototype (left); the exemplar model assumes that categories are represented by the specific encountered category exemplars (right). (D) Illustration of model-based parametric regressors used in fMRI analyses. The onset regressor (on/off) were modulated by trial-by-trial similarity to prototype or exemplar representations. The x-axis represents time (one unit per trial), and the y-axis indicates similarity with higher values reflecting greater similarity to the relevant representations.

#### Training procedure

Participants viewed on-screen illustrations on a laptop and were instructed to classify each into one of two families by pressing a key corresponding to category labels presented on the left and right sides of the screen. Following response, a correct/incorrect symbol indicated accuracy (Figure 1B left panel). Training included 240 trials, presented in five blocks of 48 trials each. Each block repeated the eight training stimuli six times in randomized order, with self-paced breaks between blocks. The prototypes were not shown.

#### Generalization procedure

Following training, participants underwent fMRI during two runs lasting 5-min-14-s each. They were instructed to classify new stimuli without feedback, indicating their response on two hand-held button boxes (Figure 1B right panel). Each trial included a 5-second stimulus presentation followed by a 4-second intertrial interval with a fixation cross. Each run comprised 34 trials in randomized order, including eight training exemplars, two prototypes, and 24 novel exemplars (12 per category, with four at each distance: 1-, 2-, and 3-away) that differed between runs.

### fMRI

A 3T Siemens Prisma-Fit MRI scanner acquired two 5-min-14-s functional categorization runs, a 6-minute resting-state scan, each with field maps, and T1-, T2-, and diffusion-weighted scans. Only data from the categorization runs are reported here (157 volumes per run, 2-second repetition time, 2-mm^3^ voxels). Preprocessing was performed with fMRIPrep v24.1.1. Details are provided in SM.

### Analytic plan

#### Computational models

Two similarity-based computational models, prototype and exemplar, were applied to each participant’s trial-by-trial responses during generalization following Bowman and Zeithamova (Bowman & Zeithamova, 2018). Both models assume that learners form mental representations of categories and categorize novel stimuli by comparing them to these internal representations. The models differ in how categories are represented: the prototype model posits a single abstract representation per category, while the exemplar model represents categories through experienced category members (the eight training exemplars; Figure 1C).

Model specifications are detailed in SM. For each participant, models were implemented by computing similarity between each stimulus and the two models’ respective reference, which was then converted into predicted choice probabilities. Model fit was assessed by comparing its predictions with participants’ trial-by-trial responses, providing an estimate of their reliance on prototype or exemplar representations, termed *categorization strategy*. To evaluate strategy dominance, we used Monte Carlo simulations: the actual participant’s responses were shuffled and used to fit both models 10,000 times, creating participant-specific null distributions of model fits. A strategy was classified as dominant if the observed model fit exceeded its null distribution (p <.05, one-tailed) and outperformed the alternative model (the observed difference in model fit exceeded the null distribution of differences, p <.05, two-tailed). The participant was classified as “random” if neither model fit better than chance, and as “comparable” if both models fit without differing.

#### Statistical analysis

All analyses used hierarchical linear modeling (HLM), implemented via the “fitlme” function in MATLAB, to account for the hierarchical structure of the data and to allow for unequal numbers of observations per participant (Pinheiro & Bates, 2006).

##### Behavioral analysis

Four HLMs were fit to examine training and generalization performance, each with a nested structure in which the measurement units corresponding to the key variable (block, feature distance, familiarity (whether a given stimulus was used during training), or model fit of the prototype and exemplar models) were nested within timepoint (T1, T2), and timepoint was nested within subjects. Each model included the key variable, timepoint, and their interaction as predictors, with stimuli set (Fish, Bugs, Butterflies) and sex assigned at birth (M, F) as covariates. Random intercepts and slopes were specified at the subject and subject × timepoint levels. See model formula in SM.

The key variables in the four behavioral HLMs were as follows: (1) Training accuracy across blocks. The key predictor was block, coded as a continuous variable (1–5); a positive effect indicates learning, and the interaction tests whether learning curves differ across timepoints. (2) Generalization accuracy by feature distance. The key predictor was feature distance, coded as a continuous variable (0–3); a negative effect reflects a typicality gradient. (3) Familiarity effect in generalization. The key predictor was familiarity (training vs. novel 2-away exemplars). (4) Model comparison. The key predictor was model type (prototype vs. exemplar), with model fits as the dependent variable, testing which model better explained performance at the group level.

##### fMRI analysis

###### First-level

First-level models included three regressors, one modeling all trial onsets accounting for general categorization task performance and two orthogonalized parametric regressors (Figure 1D) capturing trial-by-trial similarity based on the prototype and exemplar models. These parametric regressors reflected variation in similarity between stimuli and their category reference across all trials. Regions correlating with these regressors were interpreted as encoding the corresponding representation (Bowman & Zeithamova, 2018; Mack, Preston, & Love, 2013). First level models also included standard covariates of no interest: six parameters obtained by rigid body correction of head motion, white matter and cerebrospinal fluid signal. The two runs were averaged after first-level analysis.

###### Group-level

For all neuroimaging analyses, sex and mean frame-wise displacement (FD) were included as covariates; stimulus set was not included because it was not significant in the behavioral models. Two analyses were conducted, a regions of interest (ROI) analysis in native space and a whole-brain analysis in normalized space. For ROI analysis, three bilateral ROIs (VMPFC, anterior and posterior hippocampus divided at its midpoint) were defined through Desikan–Killiany and subcortical atlases in each subject’s native space using FreeSurfer. For each participant, separately for prototype and exemplar regressors, beta estimates were averaged across voxels within each ROI and analyzed using HLMs with timepoint nested within subjects and intercept allowed to vary across individuals. We tested whether the intercept exceeded zero, with Bonferroni correction for the three ROIs (p<.017).

A voxel-wise whole-brain analysis was performed using the same HLM as ROI analysis to identify brain regions tracking prototype or exemplar representations in the gray matter. Multiple comparisons were corrected using the random field theory (RFT) (voxel-wise p<.005, cluster-wise p<.05).

##### Association analysis with behavior

To evaluate the degree to which behavioral and neural correlates during generalization predict real-world functioning in autism, we performed two association analyses using HLMs. (1) Generalization accuracy and adaptive functioning as measured by ABAS-GAC. Stimuli set and sex were included as covariates. 2) Brain–behavior relationships were assessed within significant ROIs and clusters to test if neural fit was associated ABAS-GAC. Significance was set at a Bonferroni-corrected level (p<.01). For both analyses, we further examined whether these associations varied by autistic traits, as measured by total SRS score. Participants were divided into two groups based on the clinically meaningful cutoff T-score of 70 (average across T1 and T2; T-score>70 = high autism traits; T≤70 = moderate-low autism traits). See SM for detailed model specifications.

## Results

### Performance summary

To evaluate whether autistic adolescents could acquire and generalize category structures, we first examined task performance across timepoints. Among participants who completed scanning (T1: n=61, T2: n=57), those with poor data quality were first excluded (T1: n=3, T2: n=7, see SM for details). At training, eight participants did not meet learning criterion of >.55 final-block accuracy at each timepoint, with five overlapping across timepoints. After excluding these participants, the final sample included 50 (82%) participants at T1 and 42 (74%) at T2, with N=39 at both T1 and T2. Percentage of final-block correct responses averaged 80.1% (SD=13.6%) at T1 and 84.0% (SD=12.9%) at T2. During generalization, mean percentage of correct responses to novel items were 74.2% (SD=12.5%) at T1 and 78.0% (SD=9.2%) at T2. See the full distribution in Figure S3.

### Learning and generalization of category structure

During training, accuracy increased linearly as the training progressed (t(448)=11.2, p<.0001, Figure 2A), indicating robust learning. The block × timepoint interaction was not significant (p=.43), suggesting similar learning curves across timepoints. Mean accuracy at T2 was significantly higher than at T1 (t(448)=3.1, p=.002), consistent with greater experience with the task. During generalization, accuracy declined linearly as similarity to the prototypes decreased (t(361)=−11.4, p<.0001), showing a typicality gradient consistent with encoding category similarity structure (Figure 2B). Neither the main effect of timepoint nor interaction of typicality with timepoint was significant (ps≥.16), indicating similar typicality gradients at both T1 and T2. Accuracy did not differ between familiar and novel 2-away items (t(177)=1.0, p=.31), suggesting that familiarity for individual exemplars did not aid categorization performance. Neither interaction nor main effect of timepoint was significant (ps≥.2). These results demonstrate that the majority of participants successfully learned the category structure and applied what they learned to new exemplars.

**Figure 2.**
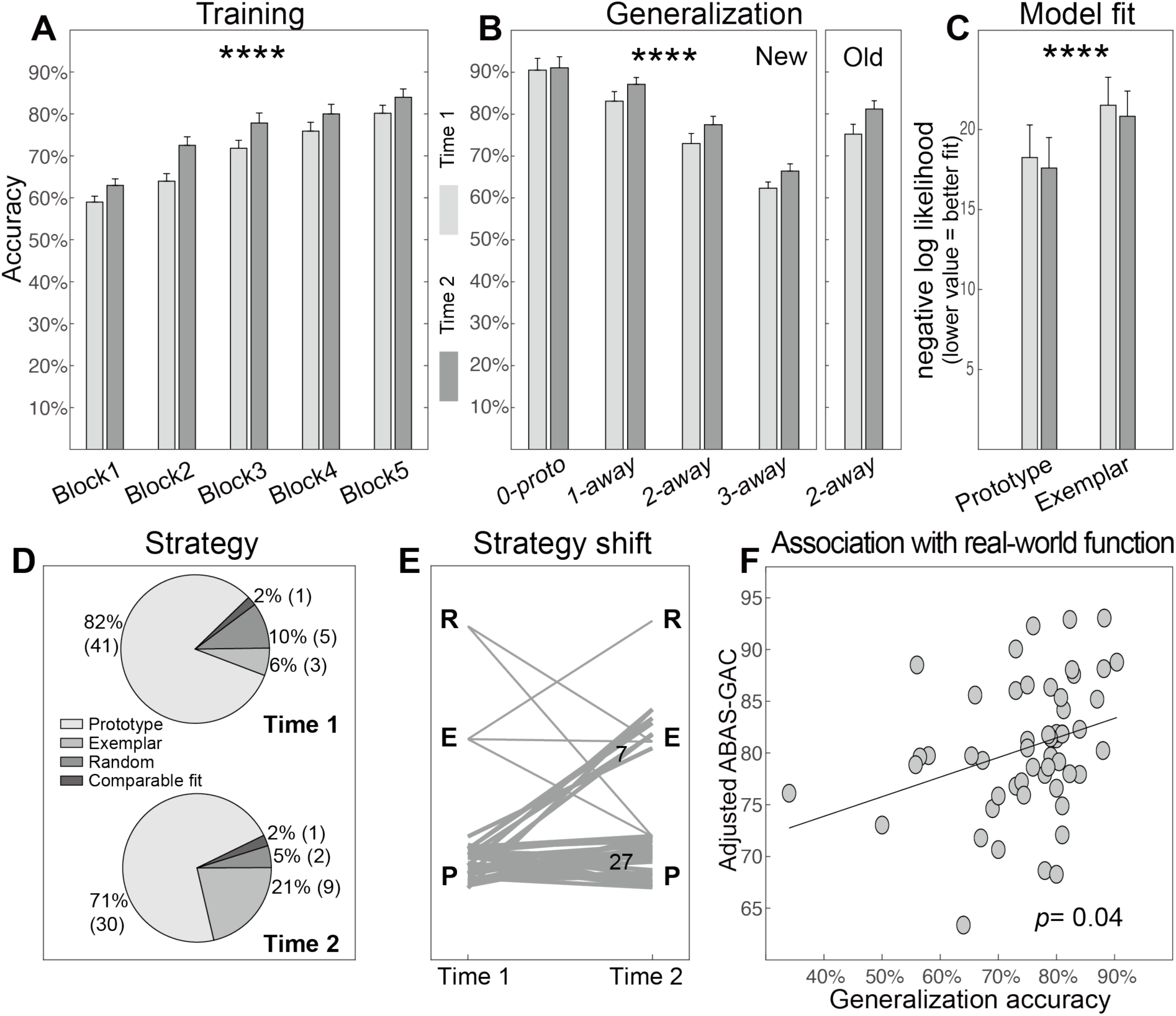
Behavioral results. (A) Training accuracy (mean % correct) by block for timepoint 1 (light bar) and timepoint 2 (dark bar). (B) Generalization accuracy (mean % correct) for new stimuli at each feature distance (0, 1, 2, 3) and for old (training) items at 2-away for each time-point. Asterisks indicate significant linear effect of block in A and feature-away condition in B (p < .0001, hierarchical linear modeling). (C) Mean model fit for the prototype and exemplar models, for each timepoint. Asterisks indicate significantly better prototype than exemplar model fits (p < .0001, hierarchical linear modeling). In panels A–C, error bars represent standard error of the mean. (D) Pie chart showing the distribution of categorization strategies across timepoints. (E) Spaghetti plot representing shift in categorization strategy from timepoint 1 to timepoint 2 among the 39 participants who have data available at both timepoints (n=27 had prototype strategy at both timepoints; n=7 shifted from a prototype strategy to an exemplar strategy; participants classified as comparable-fit were grouped with prototype). (F) Positive association between mean generalization accuracy and parent-reported adaptive functioning measured by the Adaptive Behavior Assessment System – General Adaptive Composite (ABAS-GAC) using hierarchical linear modeling. Higher GAC scores indicate better adaptive functioning.

### Dominance of prototype strategy in generalization performance

To identify the representational processes underlying this generalization, we next compared the fits of prototype and exemplar models. At the group level, the prototype model outperformed the exemplar model (t(177)=4.8, p<.0001, Figure 2C). Neither interaction nor main effect of timepoint was significant (ps≥.98), indicating that overall prototype-dominance was consistent across timepoints. At the individual level, proportion of participants by strategy is presented in Figure 2D. The majority of the participants were classified as prototype dominant: 82% (n=41/50) at T1 and 71% (n=30/42) at T2. Due to imbalanced sample size across strategy groups (prototype, exemplar, and random), no group comparison was conducted; nevertheless, we plotted accuracy, demographic and behavioral measures by strategy groups, no clear patterns were apparent (Figure S3). Strategy dominance was stable in the majority of participants who completed both assessments, with 69% (27/39) prototype-dominant and 1 exemplar-dominant at both timepoints. Of the 11 who were unstable, most (7) switched from prototype at T1 to exemplar at T2, all other switches occurred in only one case each (Figure 2E). The 27 participants with stable prototype dominance had numerically higher average generalization accuracy and lower autistic traits relative to others (Table S2).

### Generalization accuracy is associated with adaptive functioning

Mean generalization accuracy was positively associated with ABAS-GAC (t(86)=2.1, p=.04) such that participants who performed better on the generalization task also showed greater functional independence in daily life (Figure 2F). This relationship did not differ between subgroups defined by autistic traits (p=.36; high=SRS>70, N=28; moderate-low= SRS≤70, N=25).

### ROI analysis: VMPFC supports prototype representation

After establishing behavioral evidence for prototype-based generalization, we next examined its neural correlates using model-based fMRI. Significant prototype neural fit was found in the VMPFC (t(66)=2.7, p=.004), but only at a weak trend in the anterior hippocampus (t(66)=1.3, p=.10; Figure 3A). VMPFC also showed stronger prototype than exemplar neural fit (t(136)=2.7, p=.01). In the posterior hippocampus, exemplar neural fit was not significant (p=.49; Figure 3B). For each ROI, main effect of timepoint did not differ for either prototype or exemplar neural fits (ps>0.2), indicating similar neural engagement across timepoints.

**Figure 3.**
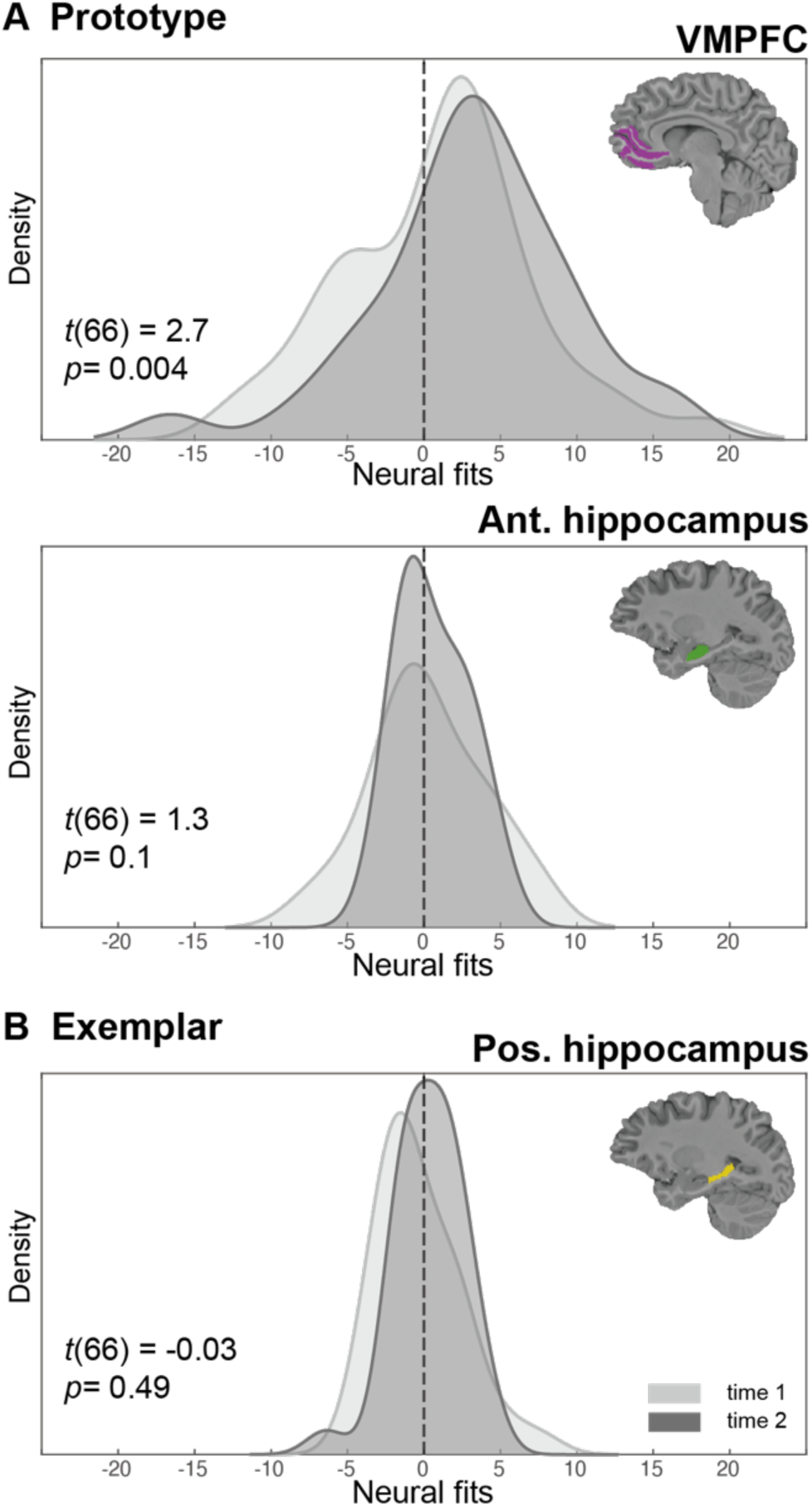
Distribution plots of prototype and exemplar neural model fits (beta) in the three bilateral regions of interest, ventromedial prefrontal cortex (VMPFC), anterior (ant) and posterior (pos) hippocampus. Light gray = time 1, Dark gray = time 2. T and p values are from intercept tests in hierarchical linear modeling.

### Whole-brain analysis: Representation correlates in occipital, parietal, and frontal cortices

To determine whether prototype representations extended beyond the a priori regions of interest, we conducted a whole-brain analysis. Four clusters showed significant prototype-related activations, posterior to anterior, in the right lateral occipital cortex (rLOC), two clusters in inferior parietal lobule (IPL) extending from supramarginal gyrus into angular gyrus in both hemispheres, and finally in the right frontal pole (rFP, Figure 4A). Significant exemplar neural fit was observed in only one cluster, in the bilateral cuneus cortex (Figure 4B). See details of these clusters in Table 2. Only rFP exhibited main effect of timepoint, with higher prototype-related activation at T2 than T1 after Bonferroni correction (t(66)=3.5, p=.001).

**Figure 4.**
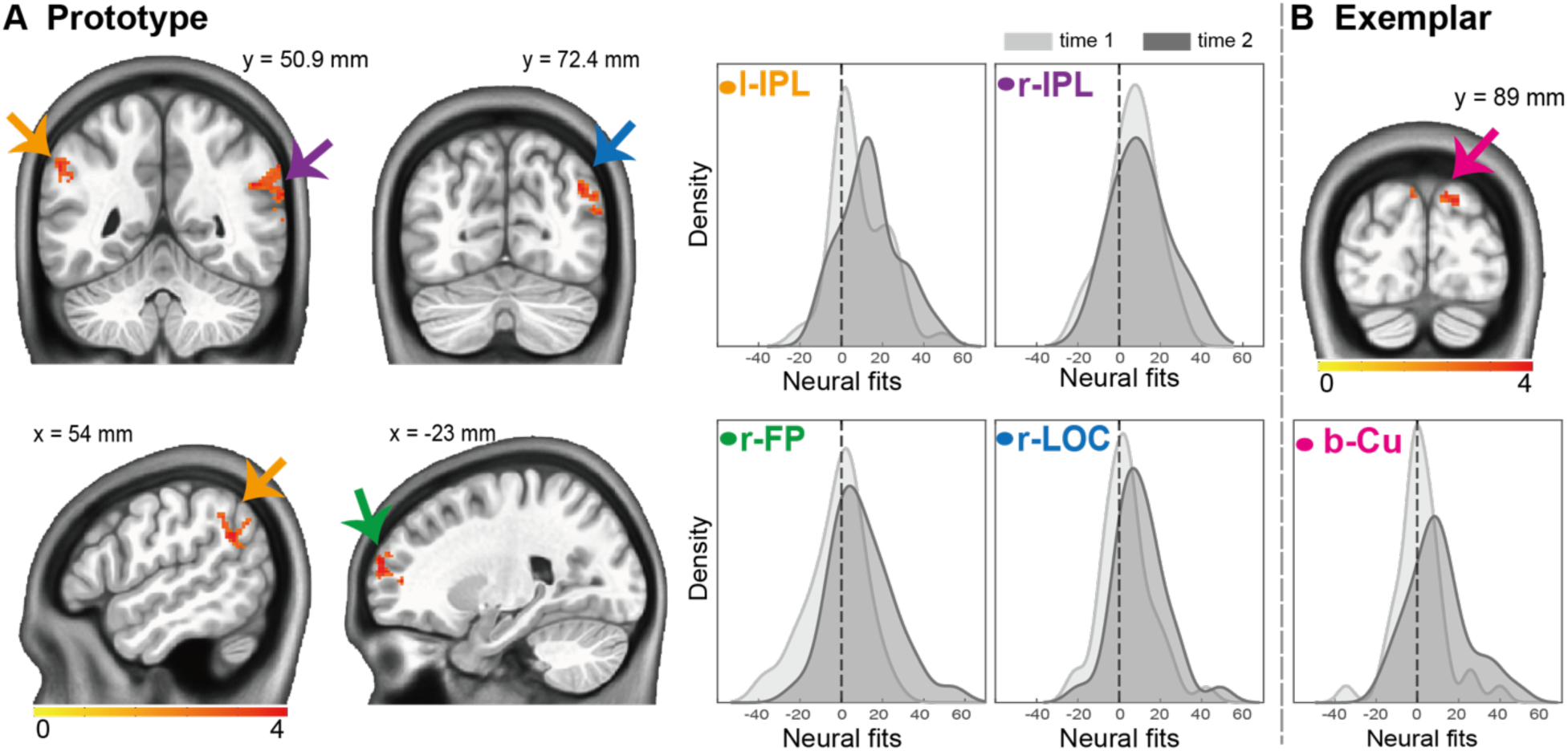
Whole-brain results for prototype and exemplar neural fits. Clusters surviving correction for multiple comparisons based on random field theory (voxel-wise p < .005, cluster-wise p < .05) are displayed with t values shown within each cluster using a yellow–red color scale for prototype (A) and exemplar (B) regressor from the hierarchical linear modeling. Images are shown in MNI space. Arrows indicate the four significant clusters: yellow = left inferior parietal lobule (l-IPL), purple = right inferior parietal lobule (r-IPL), blue = right lateral occipital cortex (r-LOC), green = right frontal pole (r-FP), pink = bilateral cuneus (b-Cu). Distribution plots show neural fit (beta) values for each cluster (light = time 1, dark = time 2).

**Table 2.** Regions significantly tracking prototype and exemplar model neural fit.

| Prototype |  | Region (Harvard-Oxford) |  |  | MNI peak coordinate |  |  |  |  |
| --- | --- | --- | --- | --- | --- | --- | --- | --- | --- |
|  |  |  |  |  | Size<br>(voxels) | z<br>value | X | Y | Z |
| r-IPL | R | 46% Supramarginal Gyrus | 40% Angular Gyrus | 5% Middle Temporal Gyrus | 470 | 4.4 | -52. | +48 | +29 |
| l-IPL | L | 39% Supramarginal Gyrus | 35% Angular Gyrus | 8% Lateral Occipital Cortex | 194 | 4.6 | +52 | +54 | +29 |
| r-FP | R | 98% Frontal Pole | - | - | 107 | 4.1 | -24 | -62 | +15 |
| r-LOC | R | 100% Lateral Occipital Cortex | - | - | 105 | 4.1 | -44 | +70 | +31 |
| Exemplar |  |  |  |  |  |  |  |  |  |
| Cuneus | Bi | 46% Cuneal Cortex | 44% Occipital Pole | 6% Precuneus Cortex | 79 | 4.3 | -16 | +90 | +35 |

### Autistic traits modulate the prototype neural fit–behavior relationship

Given the overall better fit of the prototype model, brain-behavior relationship analyses focused on prototype-related neural fits. The association between prototype-related neural fits and adaptive functioning measured by ABAS-GAC was not significant in the VMPFC and the four clusters observed with whole brain analysis (ps>.1, Figure S4). However, the interaction with autistic traits measured by total SRS scores was significant in the left IPL (t(63)=3.3, p=.001), such that higher prototype neural fit was associated with better adaptive functioning in the high-trait group (t(32)=3.2, p=.003) but not in the moderate-low group (p=.94; Figure 5; see Table S1 for demographic information). This interaction was not significant in VMPFC or the other three clusters.

**Figure 5.**
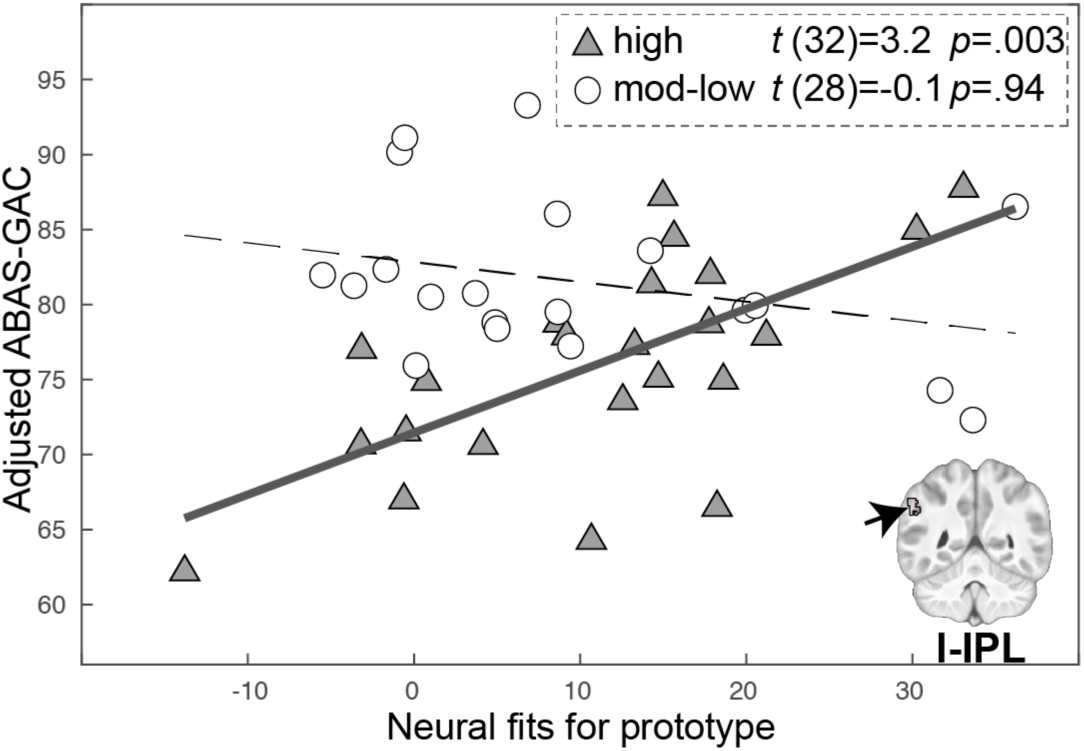
Association between neural prototype model fit in left inferior parietal lobule (l-IPL) and parent-reported adaptive functioning depends on level of autistic traits. Positive association between neural prototype model fit in l-IPL and the adaptive functioning measured by the Adaptive Behavior Assessment System – General Adaptive Composite (ABAS-GAC) in participants with high autistic traits (p = .003; corrected p < 0.05); this association was not significant in participants with moderate-low autistic traits (circles). Higher ABAS-GAC scores indicate better adaptive functioning.

## Discussion

The present study aimed to elucidate representational bases of concept learning, its stability across timepoints, and its relationship to real-world adaptive functioning in autistic youth. Prototype and exemplar models were applied to the generalization phase of a perceptual category learning task. Model-based parametric regressors probed neural correlates in the brain. The majority of autistic adolescents successfully acquired category structure and transferred this knowledge to novel stimuli. Contrary to our hypothesis, most participants’ categorization predominantly relied on prototype rather than exemplar-specific information, with consistency at a second time-point nine months later. Model-based fMRI confirmed prototype representation in the VMPFC but not anterior hippocampus, and revealed involvement of right frontal pole, bilateral IPL and right lateral occipital cortex. Exemplar representation was supported by bilateral cuneus, and not posterior hippocampus as hypothesized. Importantly, generalization predicted real-world parent-reported adaptive behavior and higher prototype-related activation in left IPL was associated with better adaptive functioning in the autistic participants with higher autistic traits. Together, these results elucidate heterogeneity in abstract representations underlying concept learning and relevance to adaptive functioning in autism.

Our results provide the first evidence challenging the prevailing view that concept learning in autism relies primarily on hyper-specific perceptual processing. Computational modelling revealed that approximately three quarters of autistic adolescents relied on prototype representations rather than exemplar-specific information during generalization of category structure. Additionally, participants’ categorization performance showed no advantage for familiar over novel items, suggesting that exemplar-specific information was not contributing to categorization. Moreover, 70% of prototype-dominant participants maintained the strategy preference over nine months, suggesting a cognitive trait rather than a transient state. Youth with consistent prototype dominance appeared to have higher generalization and lower autistic traits than the unstable subgroup (Table S2), an observation that should be statistically confirmed with more balanced sample sizes in future work. It is important to note that prior computational studies in TD adults show that task characteristics can bias categorization strategy, with higher similarity among exemplars (Bowman & Zeithamova, 2020) and feedback-based training (Houser, Resnick, & Zeithamova, 2024) promoting prototype reliance. Both attributes characterized our task which could have encouraged prototype reliance, suggesting that autistic youth conform to commonly observed representational tendencies.

Neuroimaging results largely confirmed engagement of theorized neural correlates of concept learning. First, as hypothesized, prototype correlates were observed in VMPFC, which supports integration across experiences into abstract knowledge (Zeithamova et al., 2019). The right frontal pole also tracked prototype information. This region supports both rule-based and similarity-based concept learning (Li & Deng, 2025; Paniukov & Davis, 2018; Reber, Stark, & Squire, 1998; Sherman et al., 2023) specifically tracking prototype distance (Wu et al., 2020) and episodic memory retrieval (Rugg, Otten, & Henson, 2002). Notably, it was the only region sensitive to timepoint, higher at T2 than T1, suggesting task familiarity may promote engagement of those processes. Second, prototype correlates were observed in bilateral IPL, a region frequently implicated in perceptual category representation in both human and macaque studies (Fera et al., 2005; Freedman & Assad, 2016; Liu et al., 2025; Smith & Grossman, 2008; Zeithamova, Maddox, & Schnyer, 2008). Although prior studies have associated different subregions with prototype and exemplar tracking, both models suggest IPL representations may facilitate generalization to novel items through perceptual similarity–based coding (Blank & Bayer, 2022; Mack et al., 2013; Zeithamova et al., 2008). Third, extrastriate visual cortex was engaged by both representations, in right lateral occipital cortex for prototype and bilateral cuneus for exemplar representation. In past model-based studies with TD adults, occipital engagement appears to vary, observed in lateral occipital cortex for exemplar correlates (Bowman & Zeithamova, 2018; Mack et al., 2013) and in superior occipital cortex for prototype correlates (Bowman & Zeithamova, 2018). Similar to Bowman et al (Bowman & Zeithamova, 2018), whole brain analysis revealed no additional significant exemplar correlates, suggesting that overall, exemplar information was more weakly represented than prototype. This aligns with the overall prototype dominance observed in behavior. Finally, hypothesized hippocampal engagement of anterior prototype correlates was marginally significant and posterior exemplar correlates were not observed. These weak representations may reflect higher heterogeneity, but future work is needed to clarify whether it is specific to autism or adolescence in general.

Our results provide new insights into heterogeneity in concept learning that could be harnessed for intervention efforts. First, we observed three subgroups, a small minority who failed to acquire category structure, one-third who generalized without consistent representational reliance, and a consistent prototype-dominant majority. Representation dominance was not distinguished by any demographic or performance variables (Figure S3). These findings inform stratified intervention approaches: those without consistent representational reliance may benefit from explicit instruction on abstraction, to help “see” the commonalities across experiences, whereas individuals unable to learn category structure may require alternate training regimens. Second, the clinical relevance of concept learning is bolstered by the observed links to adaptive functioning. While theorized, our results provide the first empirical support for the association between generalization success on a laboratory task and effective daily application of social and independent living skills as reported by parents. The ability to form generalized representations may be particularly important for those skills, as higher prototype-related activation in the left IPL was associated with better adaptive functioning specifically in participants with higher autistic traits, highlighting a potential neural treatment target. The left IPL contributes to integration of sensory and motor information in the service of symbolic and mnemonic processing (Igelström & Graziano, 2017), and its response to intervention should be evaluated in future work.

Interpretation of the current findings is constrained by the following factors: First, similar to national estimates, psychotropic medication use was highly prevalent in our sample (Spencer et al., 2013). While stimulants were withheld during scanning, other medications could not be without risking clinical care. Second, generalizability is limited to autistic adolescents without intellectual disability as inclusion was restricted to IQ>80. Third, while use of a longitudinal design enhanced reliability and statistical power, model-based fMRI compounded challenges of data retention as strict behavioral quality control was necessary to ensure sufficient trials for modelling in addition to head motion criteria.

In sum, our findings address open questions in current understanding of concept learning in autism. Specifically, they reveal capacity for abstract processing in the majority of autistic youth and provide some empirical support for theorized neural underpinnings and association with real-world adaptive behavior. Future research should investigate its potential as a treatment target and determine the extent to which abstract representations are plastic, with implications for clinical translation.

## Acknowledgments

We would like to thank Jordan Linde and Katie Flaharty, and Xiaozhen You for generating experimental materials, Zixuan Zhao and Gang Chen for statistical advice, and participants and their families for their time. This work was supported by the District of Columbia Intellectual and Developmental Disabilities Research Center (DC-IDDRC) Award P50HD105328 by NICHD (PI: William Gaillard).

