## Supplementary material for "From cognitive abstraction to adaptive behavior: neural bases of concept learning in autistic adolescents": SM

### SUPPLEMENTARY MATERIALS

#### Methods

##### Medication and comorbidities

Of the 64 participants enrolled, 51 participants (78%) were taking at least 1 psychotropic medication, with 37 participants (57%) on stimulant medication, which was withheld for at least 24 hours before fMRI. Parents reported the following lifetime comorbidities (without distinguishing between past and current): 52 with ADHD, 49 with anxiety, 29 with depression, 14 with learning disabilities, 7 with adjustment problems, 9 with disruptive behavior disorders, 5 with conduct disorder, 3 with eating disorders, 2 with trauma-related disorders, 1 with psychosis, and 7 with other conditions not specifically queried. Three participants were reported to have no comorbidities.

##### Behavioral assessments

*Adaptive Behavior Assessment System-II (ABAS-II).* Parents completed the school-age ABAS-II, a widely used standardized questionnaire assessing adaptive functioning across daily life settings, including home, school, and the community. It assesses nine domains—communication, functional academics, self-direction, leisure, social, community use, home living, health and safety, and self-care—providing a Global Adaptive Composite (ABAS-GAC) standard T-score (M = 100, SD = 15). Higher scores indicate better adaptive functioning. ABAS-GAC scores remained stable across assessments ($r=0.82$, $p<0.001$, Figure S2).

*The Social Responsiveness Scale-II (SRS-II).* Parents completed the SRS-II, a 65-item standardized questionnaire assessing social communication and interaction and restricted and repetitive patterns of behavior associated with autistic traits. Ratings are expressed as a total T-scores (M=50, SD=10), with higher scores reflecting greater social difficulty and more severe autistic symptoms. The total SRS scores remained stable across timepoints; therefore, we averaged the scores to provide an index of autism severity ($r=0.81$, $p<0.001$, Figure S2).

#### MRI Acquisition

Scanning was completed on a 3T Siemens Prisma-Fit MRI scanner at Georgetown University using a 64-channel head coil and foam pads for head stabilization. The scan sequence included two functional categorization runs, a 6-minute resting-state scan, field maps for each functional run, and T1-, T2-, and diffusion-weighted scans. Functional volumes, consisting of 72 slices (1.8 mm thickness), were acquired in an ascending interleaved order using a multiband gradient-echo pulse sequence: TR = 2000 ms, TE = 29 ms, flip angle = 90°, FOV = 208 × 208 mm, voxel size = 2.0 × 2.0 mm, multiband acceleration factor = 3, and 157 volumes per categorization run. A B0 field map was generated using a double-echo, gradient-recalled echo sequence: TR = 697 ms, TE 1 = 4.92ms, TE 2 = 7.38 ms. High-resolution T1-weighted anatomical images were acquired using an MPRAGE sequence: TR = 1900 ms, TE = 2.52 ms, TI = 900 ms, flip angle = 9°, 176 axial slices with 1.0 mm thickness, FOV = 256 × 256 mm, voxel size = 1.0 × 1.0 mm; GRAPPA factor = 2. Given the susceptibility of the orbitofrontal region to signal loss from magnetic field inhomogeneities, we addressed this issue in two ways: 1) functional volumes were oriented 15° off the anterior commissure-posterior commissure line to reduce orbitofrontal signal dropout ; 2) voxels with BOLD signal intensities more than 2 SD below the brain-wide mean were excluded.

#### MRI Preprocessing

Raw DICOM images were converted to BIDS format using dcm2bids v3.1.1. Preprocessing was performed with fMRIPrep v24.1.1, including slice-timing correction, motion correction, and fieldmap-based distortion correction. Functional images were co-registered to the T1-weighted image using boundary-based registration (native space outputs) and normalized to MNI152NLin2009cAsym space (MNI space outputs) (1). Preprocessed images were then smoothed (4 mm FWHM), intensity-normalized, and high-pass filtered (90 s) prior to analysis, using FEAT in FSL. Following preprocessing, the two categorization runs underwent first-level analysis separately and were then averaged using a fixed-effects model in FEAT.

**Quality control**

For behavioral analyses, exclusion criteria included: 1) missed >33% trials per run; 2) response bias (>75% same-button responses) with mean generalization accuracy <.55 per run; 3) task disengagement based on self-report key-press pattern (e.g., both keys pressed, key press held through the trial or 0 reaction time; 4) failure to learn defined as training accuracy <.55 in the final block. Neuroimaging exclusions included: (1) gross anatomical abnormalities; (2) preprocessing failure; (3) excessive head motion (>20% of volumes with framewise displacement (FD)>1 mm); (4) high prototype–exemplar regressor correlation ($r > .95$), introducing collinearity in first-level analysis; (5) absence of occipital activation ($z<1$, Desikan–Killiany-defined) with mean generalization accuracy <.55, suggesting task disengagement.

**Computational models**

*Prototype model.* For each participant, the model was implemented by computing similarity between each stimulus and the two prototypes. Physical distance to each prototype was computed in feature space for each stimulus, weighted by attention parameters across the eight dimensions. These weights reflected attention allocated to individual features. Similarity was then computed using an exponential decay function, assuming that perceptual similarity declines with increasing distance:

$$Sim_A(x) = \exp\left[-c\sum_{i=1}^{8}\left(w_i\left|x_i - \text{proto}_{Ai}\right|^r\right)^{1/r}\right] \tag{1}$$

Sim$_A$ $(x)$ denotes similarity between stimulus $x$ and Prototype A; $x_i$ and $\text{proto}_{Ai}$ are 1×8 binary vectors representing feature codes (Figure 1A); $w_i$ is a 1×8 vector of attention weights, constrained to sum to 1; $c$ is a sensitivity parameter, which controls similarity decay with distance and was constrained to be 0–100; and $r$ was fixed at 1, implementing a city-block metric appropriate for binary features.

This yielded two similarity values per trial (SimA, SimB), which were converted into predicted choice probabilities (e.g., ProbA = SimA / (SimA + SimB)). To assess alignment between the participant's response and the model prediction, we assigned the predicted choice probability of the chosen category as the alignment score (i.e., per-trial likelihood). For example, choosing A with ProbA = 0.7 yields an alignment score of 0.7; choosing B gives 0.3. Overall model fit was calculated by summing the negative log-transformed alignment scores (i.e., negative log-likelihoods) across trials, where lower scores indicate better correspondence between model and behavior.

_Exemplar model_. The exemplar model followed the above procedure to calculate similarity but used the eight training exemplar (four per category) as reference points. For each stimulus, similarity to each exemplar was computed, and category-level similarity (SimA, SimB) was obtained by summing similarities within each category. Subsequent steps matched those of the prototype model.

$$Sim_A(x) = \sum_{y \in A} \exp\left[-c \sum_{i=1}^{8} (w_i |x_i - y_i|^r)^{1/r}\right] \tag{2}$$

$y$ is a training exemplar from category A.

All 68 trials were included in model fitting. Training and prototype trials were included to enable fitting both prototype and exemplar models. Missed trials were excluded. Attention weights and the sensitivity parameter were optimized so that the model predictions are as close as possible to the actual responses, separately for each model and participant using maximum likelihood estimation via MATLAB's fmincon.

Per-trial summed similarity across categories (SimA + SimB) was used to modulate the onset regressor (on/off) to create parametric regressors for the fMRI analyses, as it indexes how strongly a stimulus matches the overall category representations and has been used in model-based studies to localize brain regions encoding such representations (2,3). Non-response trials were excluded from model fit calculation but retained for regressors, as the per-trial summed similarity depended solely on stimulus features once parameters were determined.

**Statistical analysis**

*Behavioral analysis: hierarchical linear models*

This model illustrates analysis for Training accuracy. Accuracy for block k, timepoint j, subject i was modeled as:

$$\begin{aligned} \mathrm{ACC}_{ijk} = {} & \mathrm{intercept} + \beta_1(\mathrm{block}) + \beta_2(\mathrm{timepoint}) + \beta_3(\mathrm{block} \times \mathrm{timepoint}) \\ & + \beta_{\ldots}(\mathrm{covariates}) + d_i + d_{ij} + e_{ijk} \end{aligned} \tag{3}$$

Here, the intercept and β terms are fixed effects, $d_i$ is the subject-level random effect, $d_{ij}$ is the measurement-level random effect nested within subjects, and $e_{ijk}$ is the residual. The intercept and $\beta_1$ were allowed to vary across sessions and individuals.

*Association analysis: hierarchical linear models*

(1) Generalization accuracy and adaptive functioning as measured by ABAS-GAC. Association analyses using longitudinal data face the challenge that the between- and within-subject correlations are mixed. While the between-subject effect evaluated the overall effect of accuracy on ABAS-GAC, as measured by the association of the mean accuracy, the within-subject effect reflected how changes in accuracy across timepoints affect ABAS-GAC within a subject. To separate these, we decomposed the between-subject from the within-subject accuracy effect using the methods of Neuhaus (4). As this study pooled two timepoints to boost power rather than test for longitudinal change, the between-subject effect ($\beta_B$) was the primary focus. Task version and sex were included as covariates.

$$\mathrm{ABAS\_GAC_{ij}} = \mathrm{intercept}\ \overbrace{+\beta_B \overline{\mathrm{acc}}_\mathrm{i}}^{\text{between-subject}}\ \overbrace{+\ \beta_W\left(\mathrm{acc_{ij}} - \overline{\mathrm{acc}}_\mathrm{i}\right)}^{\text{within-subject}} + \beta_{\ldots}(\mathrm{covariates}) + d_\mathrm{i} + e_\mathrm{ij} \quad (4)$$

$\overline{\mathrm{acc}}_i$ is subject i's mean accuracy; the intercept and $\beta_w$ were allowed to vary across individuals.

2) Brain–behavior relationships were assessed within significant ROIs and clusters to test if neural fit was associated ABAS-GAC. Mean FD and sex were included as covariates. The effect of brain (neural fit) was decomposed into between- and within-subject components. We further examined whether the between-subject brain effect varied by autistic trait severity. Significant interactions were followed by post hoc group-specific analyses.

$$\text{ABAS\_GAC}_{ij} = \text{intercept} \overbrace{+\beta_B \overline{\text{brain}}_i}^{\text{between-subject}} + \overbrace{\beta_W\left(\text{brain}_{ij} - \overline{\text{brain}}_i\right)}^{\text{within-subject}} + \beta_2(\text{symptom}_i) + \beta_3\left(\overline{\text{brain}}_i \times \text{symptom}_i\right) + \beta_{\dots}(\text{covariates}) + d_i + e_{ij} \quad (5)$$

### Results

#### Data quality control

Participants with poor behavioral data quality were first excluded (T1: n=3, final block: M=86.8%, SD = 22.9%; generalization: M = 44%, SD = 4%; T2: n = 7, final block: M = 66.4%, SD = 15.7%; generalization: M = 54%, SD = 14%). Next, participants who failed to learn were removed (T1: n = 8, final block: M = 47.9%, SD = 3%; generalization: M = 50%, SD = 14.4%; T2: n = 8, final block: M = 50.5%, SD = 1.9%; generalization: M = 54.4%, SD = 14.1%), including five at both timepoints. Participants with below chance generalization were retained (N=4 at T1 and N=1 at T2 with accuracy<55%). The final sample included 50 of the 61 (82%) participants who completed the scan at T1 and 42 of 57 (74%) at T2, with N=39 at both T1 and T2.

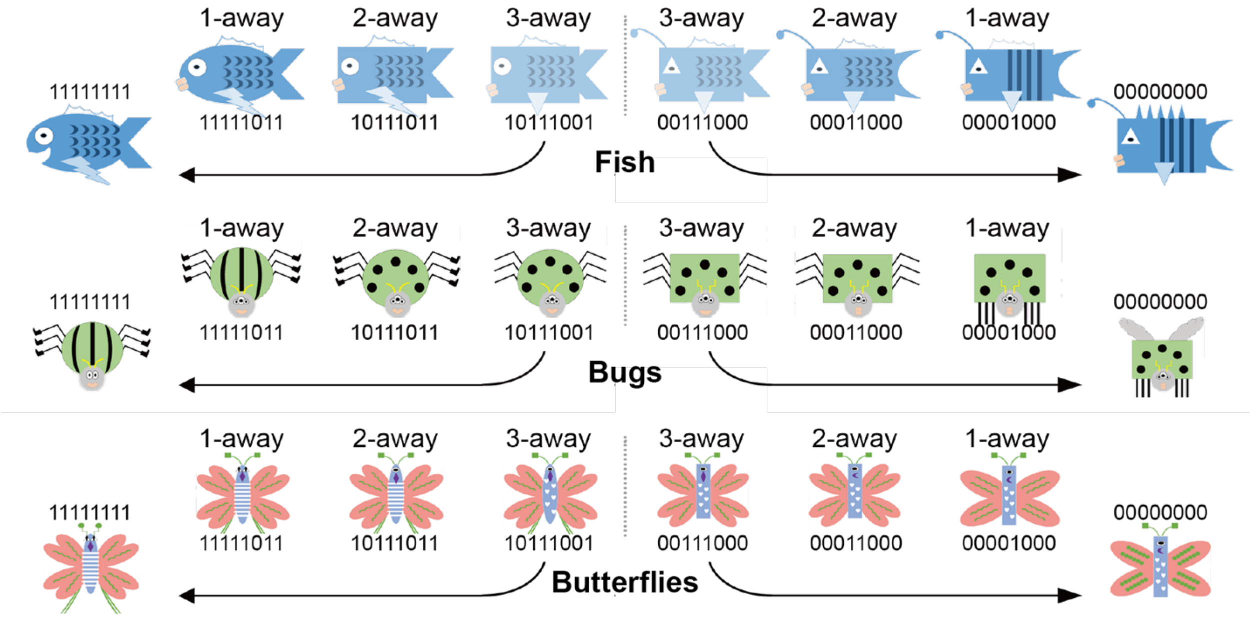


**Figure S1.** Example stimuli spanning *n* feature-distance away from the prototypes for the three cartoon animal stimuli sets used in the present study.

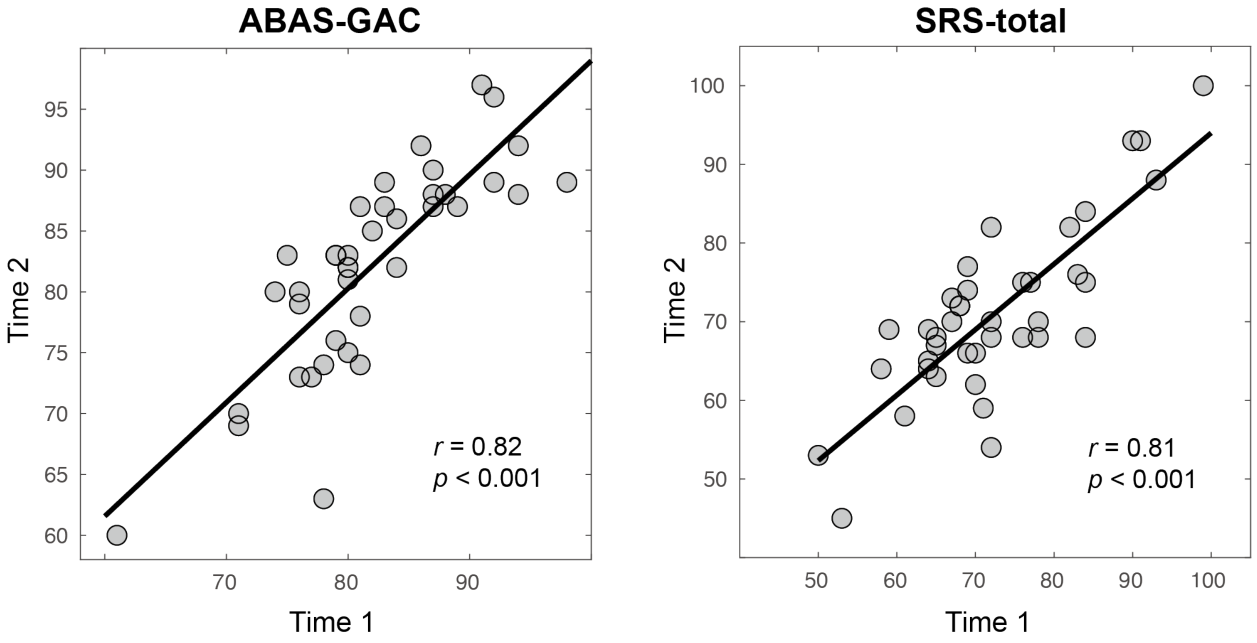


**Figure S2. correlation between timepoint 1 and timepoint 2 measures of adaptive function and autism traits.**

Adaptive functioning was assessed using the Adaptive Behavior Assessment System–General Adaptive Composite (ABAS-GAC), and autism traits were assessed using the Social Responsiveness Scale-II total score.

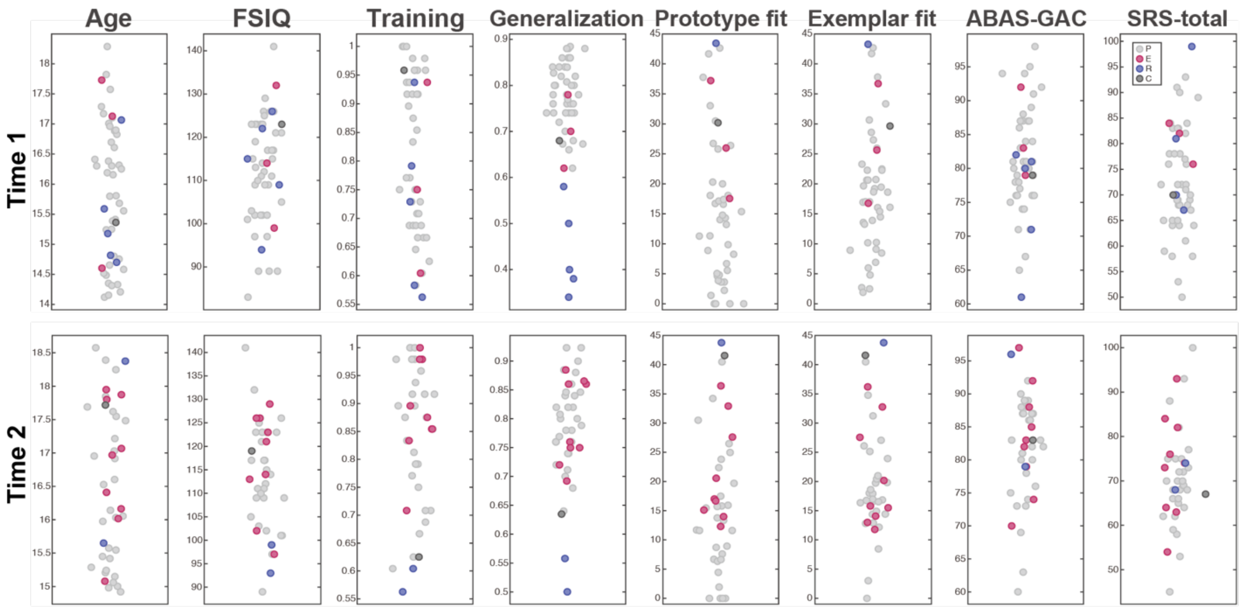


**Figure S3. Scatterplot of age, FSIQ, mean accuracy during final-block of training, mean accuracy in generalization, prototype model fit, exemplar model fit, adaptive function, and autism traits by learning strategies.**

Each dot represents a single assessment for one participant. Participants with different learning strategies are represented by different colors. P=prototype (light grey); E=exemplar (red); R=random (blue); C=comparable fit (dark grey).

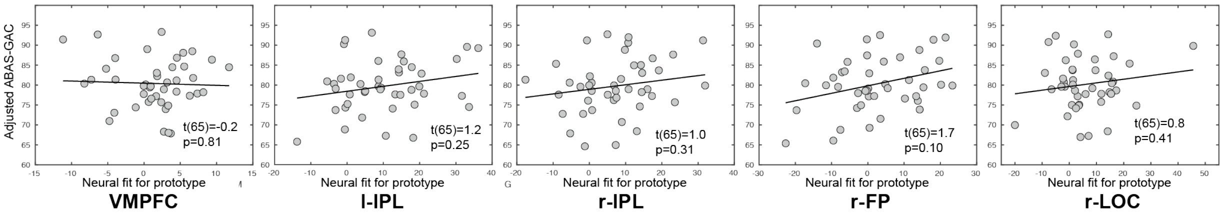


**Figure S4. Association between parent-reported adaptive functioning and prototype neural fit in the VMPFC and the four clusters observed with whole brain analysis**.

VMPFC= ventromedial prefrontal cortex; l-IPL=left inferior parietal lobule, r-IPL=right inferior parietal lobule, r-LOC=right lateral occipital cortex, r-FP=right frontal pole.

Table S1. Diagnostic and demographic characteristics of participants included in neuroimaging analysis for analysis by High autism traits and Low autism traits. The two timepoints were averaged for mean and standard deviation (SD). Autistic traits were measured by Social Responsive Scale Total score (Severe: T>70; Mild: T<70).

| **Characteristics** | Mean ± SD or n (%) | | Sig. |
|---|---|---|---|
| | **High** | **Low** | |
| #Participants | 23 | 20 | |
| Age, Years | 16.0 ± 1.2 | 15.9 ± 1.1 | p = 0.77 |
| Sex assigned at birth (female/male) | 8 / 15 | 5 / 15 | p = 0.49 |
| Self-reported Gender (female/male/non-binary) | 4 / 16 / 3 | 5 / 14 / 1 | p = 0.59 |
| Race (Nat. Am./Afr. Am./Asian/White/Others) | 1 / 2 / 1 / 19 / 0 | 0 / 2 / 0 / 17 / 1 | p = 0.57 |
| Ethnicity (Latino/Non-Latino) | 1 / 22 | 3 / 17 | p = 0.23 |
| Full Scale IQ (Standard Score) | 113.4 ± 12.9 | 115.7 ± 11.3 | p = 0.54 |
| Interval between T1 and T2, Months | 9.1 ± 0.8 | 9.0 ± 0.8 | p = 0.53 |
| Social Communication Questionnaire (Raw Score) | 17.2 ± 7.9 | 13.1 ± 6.6 | p = 0.02 |
| Social Responsiveness Scale Total Score (T score) | 79.6 ± 6.8 | 63.6 ± 5.6 | p<0.001 |
| Social Communication and Interaction (T score) | 78.5 ± 6.7 | 63.1 ± 6.0 | p<0.001 |
| Restricted Interests and Repetitive Behaviors (T score) | 80.2 ± 10.3 | 64.2 ± 6.5 | p<0.001 |
| Adaptive Behavior Assessment System Global Adaptive Composite (Standard Score) | 79.0 ± 7.9 | 84.0 ± 6.0 | p = 0.02 |
| Final-block training accuracy | 83% ± 11% | 83% ± 12% | p = 0.90 |
| Generalization accuracy | 76% ± 10% | 79% ± 8% | p = 0.34 |
| Medication | | | |
| Stimulants | 13 (57%) | 11 (55%) | |
| Non-stimulants | 6 (26%) | 1 (5%) | |
| Antianxiety / Antidepressant | 15 (65%) | 13 (65%) | |
| Anti-psychotic | 2 (9%) | 1 (5%) | |
| Other (antihistamine, Endocrine, etc.) | 5 (22%) | 6 (30%) | |

Table S2. Diagnostic and demographic characteristics in participants who have behavioral data for both two timepoints, divided into prototype-stable and others (unstable and exemplar-stable) groups. The two timepoints were averaged for mean and standard deviation (SD).

| **Characteristics** | Mean ± SD or n (%) | |
|---|---|---|
| | **Prototype-stable** | **Other** |
| #Participants | 27 | 12 |
| Age, Years | 15.6 ± 1.1 | 16.0 ± 1.1 |
| Sex assigned at birth (female/male) | 8 / 19 | 6 / 6 |
| Self-reported Gender (female/male/non-binary) | 5 / 19 / 3 | 5 / 6 / 1 |
| Race (Nat. Am./Afr. Am./Asian/White/Others) | 1 / 1 / 0 / 24 / 1 | 0 / 2 / 0 / 10 / 0 |
| Ethnicity (Latino/Non-Latino) | 4 / 23 | 0 / 12 |
| Full Scale IQ (Standard Score) | 114.4 ± 10.9 | 115.9 ± 12.1 |
| Interval between T1 and T2, Months | 9.1 ± 0.8 | 9.0 ± 0.9 |
| Social Communication Questionnaire (Raw Score) | 12.5 ± 6.9 | 17.2 ± 6.5 |
| Social Responsiveness Scale Total Score (T score) | 69.4 ± 9.3 | 76.5 ± 11.5 |
| Social Communication and Interaction (T score) | 68.4 ± 9.0 | 75.3 ± 11.5 |
| Restricted Interests and Repetitive Behaviors (T score) | 70.9 ± 11.0 | 77.3 ± 13.1 |
| Adaptive Behavior Assessment System Global Adaptive Composite (Standard Score) | 81.9 ± 6.3 | 82.1 ± 9.9 |
| Final-block training accuracy | 83% ± 11% | 84% ± 12% |
| Generalization accuracy | 79% ± 6% | 75% ± 11% |
| Medication | | |
| Stimulants | 16 (59%) | 6 (50%) |
| Non-stimulants | 4 (15%) | 1 (8%) |
| Antianxiety / Antidepressant | 14 (52%) | 11 (92%) |
| Anti-psychotic | 1 (4%) | 2 (17%) |
| Other (antihistamine, Endocrine, etc.) | 6 (22%) | 4 (33%) |
